# Restoring morphology of light sheet microscopy data based on magnetic resonance histology

**DOI:** 10.1101/2022.07.12.499779

**Authors:** Yuqi Tian, James J. Cook, G. Allan Johnson

## Abstract

The combination of cellular-resolution whole brain light sheet microscopy (LSM) images with an annotated atlas enables quantitation of cellular features in specific brain regions. However, most existing methods register LSM data with existing canonical atlases, e.g. The Allen Brain Atlas (ABA), which limits the precision of measuring regional volumes and eliminates the individual difference in brain partitions for different strains, ages, and environmental exposures. Thus, these approaches obscure valuable anatomical information. Here, we present a method to combine LSM data with magnetic resonance histology (MRH) of the same brain restoring the morphology of the LSM images to the in-skull morphology. Our pipeline which maps 3D LSM data (terabyte level per dataset) to magnetic resonance histology of the same mouse brain provides accurate registration with low displacement error in 10 hours with limited manual input. Optimization and validation, from initialization of the data, designing the quantitative loss function, optimizing the 20+ image processing variables on multiple resolution scales, and finalizing the application on full resolution data has been integrated through a structured workflow. Excellent agreement has been seen between registered data and reference data both locally and globally. This workflow has been applied to a collection of datasets with varied MRH and LSM contrast providing a routine method for streamlined registration of LSM images to MRH.

## 1. Introduction

Combining mesoscopic structural information of the brain and histology at the cytoarchitectural scale has been a focus in recent years to reveal the bridge between tissue morphological alternations and disease (Casanova et al. 2009) (Vemuri and Jack 2010) (Zhang et al. 2012), brain insult (Tuor et al. 2014) (Weishaupt et al. 2016) (Fornito, Zalesky, and Breakspear 2015) and aging (Schmitz et al. 2018) (Eylers et al. 2016). There is clear evidence that morphological disruptions underlie brain dysfunctions at both the meso- and microscopic scale; for example the corpus callosum volume reduction in autism (Egaas, Courchesne, and Saitoh 1995) (Hardan, Minshew, and Keshavan 2000) (Tepest et al. 2010) (Loomba et al. 2021) and neuronal death following ischemic insult (Weishaupt et al. 2016). Merging structural changes in specific brain regions at the mesoscale with corresponding quantitative cellular measurements at the microscopic scale will open an entirely new window into understanding the brain.

Diffusion tensor imaging (DTI) provides particularly unique insight into brain morphology and connectivity (Fornito, Zalesky, and Breakspear 2015). However, extension of DTI to more basic studies in the mouse is challenging because the mouse brain @ 435 mg is about 3000 times smaller than the human brain. Through a series of innovations, the Duke Center for in vivo Microscopy (CIVM) has extended the spatial resolution of Magnetic resonance imaging (MRI)/DTI by more than 500,000 times that of routine clinical scans in perfusion fixed post mortem specimens (e.g. MRH) (Johnson et al. 1993) (Johnson et al. 2007). Recent work has pushed the resolution of DTI to 15×15×15 μm^3^ and accelerated the acquisition with compressed sensing, which enables routine acquisition of high-resolution multidimensional dimensional whole mouse brain images (Johnson et al. 2019; Wang, Anderson, et al. 2018). These high-fidelity mesoscale MRH data now enable correlation between the MRH metrics and the tissue cytoarchitecture.

The development of tissue clearing and LSM have allowed neuroscientists to routinely image whole cleared mouse brains at cellular resolution (Erturk et al. 2012). Continued innovation in clearing (SHIELD) (Park et al. 2019) and immunohistochemistry (SWITCH) (Murray et al. 2015a) has enabled staining of varied cell types (neuron, oligodendrocyte, microglia), structural proteins (myelin) and pathologies (a-beta and tau proteins).

Merging MRH and LSM data from the same specimen will capture the best of both. MRH with DTI is a non-destructive and multi-contrast imaging tool which preserves accurate brain morphology since the scanning can be done with the brain in the skull. DTI with high angular sampling provides maps of whole brain connectivity. Multiple scalar images provide exquisite tissue contrast differentiating brain subunits. Post processing pipelines can exploit these multi-contrast images to automatically label more than 300 different sub-regions. LSM provides cellular resolution but requires the removal of the brain from the skull and tissue clearing, which induces tissue swelling. Dissection of the brain from the skull frequently results in tissue loss or tearing (Figure 1). And labeling is not always as uniform as one might hope. Mapping LSM to MRH restores the tissue geometry and allows automated labeling of the subregions in the LSM data. Finally, the most common method for labeling LSM images involves registration to the Allen Brain Atlas which has been constructed from 2D serial sections acquired at 100 μm intervals averaged from ~1600 young adult C57BL/6J mice. (Adler et al.). Mapping LSM data from another strain at a different age will obscure regional volume changes that might be important image phenotypes for the study. However, we lack efficient methods to solve this difficult multi-modal registration problem. The geometric distortion in the LSM data can exceed 40% and there is frequent tissue loss and distortion. There are profound contrast differences between the many MRH scalar images and the many potential immunohistochemistry stains. The data volumes are large approaching a terabyte for a single specimen. Our long-range goal is development of the infrastructure to support routine, comprehensive morphologic phenotyping of the mouse brain using combined MRH and LSM to map the genetic impact on cells and circuits. We describe here a workflow and pipelines that address these challenges making combined MRH/LSM of the same brain routine.

**Figure 1.**
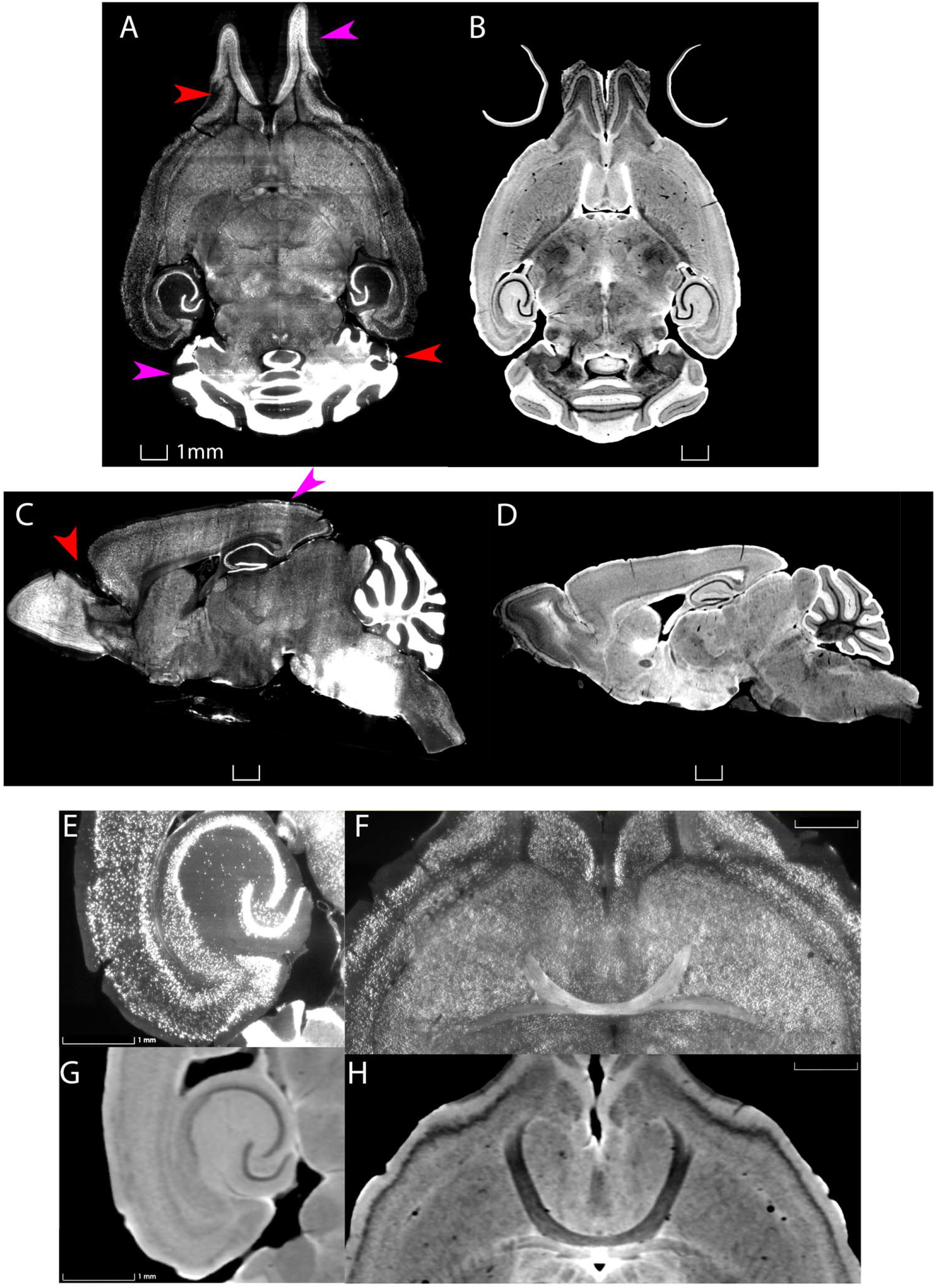
Distortion and tissue tearing in LSM compared to MRH. A comparison between LSM images of a mouse brain stained with NeuN (A,C,E,F) and a diffusion weighted MRH image of the same specimen (B,D,G,H) highlights some of the challenges and opportunities. Red arrows indicate the tissue tearing. Purple arrows indicate the swelling. Specimen 200316.

## 2. Materials and Methods

### 2.1 MRH histology

All animal procedures were conducted under guidelines approved by the Duke University Animal Use and Care Committee. Specimens were perfusion fixed using an active staining method that has been described in detail previously (Johnson et al. 2019). Warm saline to flush out blood was perfused through a catheter in the left ventricle. This was followed by a formalin/Prohance mixture titrated to reduce the spin lattice relaxation time (T1) of the tissue enabling accelerated scanning. The MRH scanning was performed on a 9.4T vertical bore magnet with a Resonance Research Inc (Billerica, Md) gradient coil yielding peak gradients up to 2500mT/m. The scanner is controlled by an Agilent console running VnmrJ 4.0. The acquisition was accelerated using compressed sensing(Johnson et al. 2019; Wang, Cofer, et al. 2018). Diffusion tensor images were acquired using a protocol described in detail in (Johnson et al. 2022). The protocol employed a Stesjkal Tanner spin echo sequence with b values of 3000 s/mm^2^, 108 angular samples spaced uniformly on the unit sphere, a compression factor of 8 x yielding a large (252 GB) 4D volume with isotropic resolution of 15 μm.

### 2.2 LSM

Following the MRH scans, the brains were removed from the skull and sent to Life Canvas Technology (https://lifecanvastech.com/) for LSM imaging. There the brains were cleared using SHIELD (Park et al. 2019) and stained using SWITCH (Murray et al. 2015b) and scanned on a selective plane illumination microscope (SPIM) yielding three channel whole brain images at 1.8×1.8 × 4.0 μm. Each of the three channels yields an isotropic volume at a different wavelength of ~ 300GB. The aggregate dataset for one specimen (MRH and 3 channels of LSM) is typically ~ 1 TB. Table 1 lists data sets that have been used to test the pipelines.

**Table 1.**
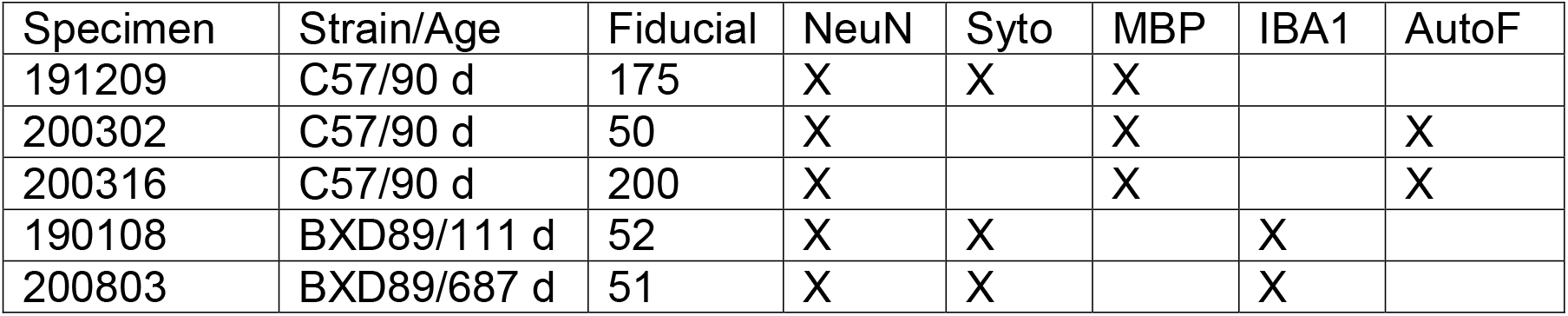
Test specimens for combined MRH/LSM registration

### 2.3 Preprocessing and initialization

Initial attempts at registration were particularly unsuccessful in cerebellum and olfactory bulb both of which are prone to significant distortion after removal from the skull (Fig.2). We developed an initialization procedure using sparse landmarks (15~20) with many concentrated in these two regions. Landmarks are placed in pairs, on both LSM and MRH. A complete landmark registration (e.g., 150~200 pairs of landmarks) takes at least a day by someone with knowledge of the neuroanatomy of the mouse brain. This preprocessing step was accelerated by establishing a set of easily identified landmarks placed in crucial points where the distortion is most significant. The landmark locations are 4 landmarks on olfactory bulbs, 2-3 landmarks on vessels on both sides between cortex and striatum, 3 landmarks on cerebellum, 2 landmarks on dentate gyrus, 2 landmarks on hippocampus and 2 landmarks on brain stem.

**Figure 2.**
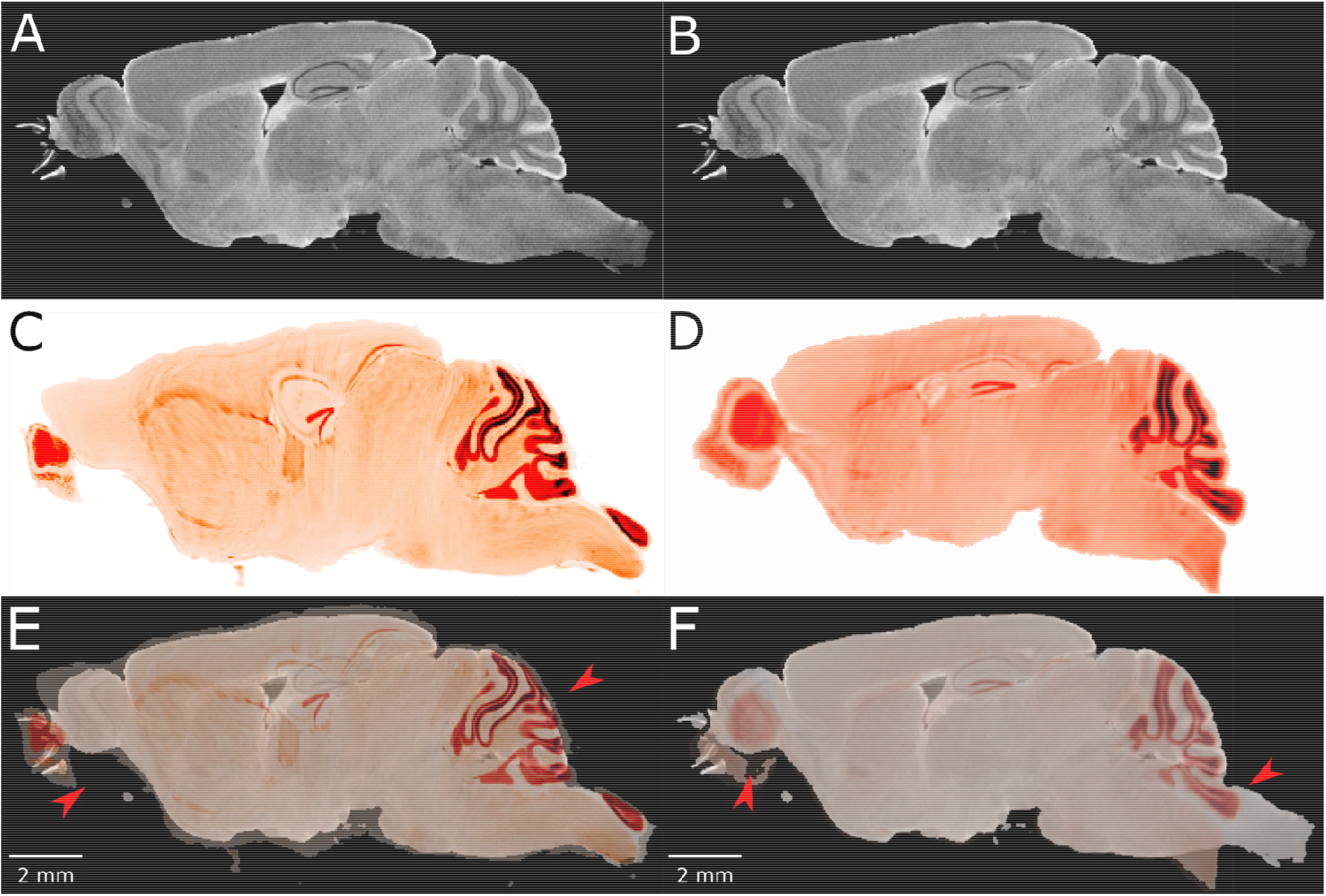
The failure of existing registration algorithms in the cerebellum and olfactory bulb. (A,B)-DWI; (C-D) NeuN image after registration; (E-F) overlaid DWI/NeuN. (Specimen 191209) The left hand column shows the result of Elastix (Klein et al. 2010) with rigid and bspline registration and default settings. The registration errors in the olfactory bulb and brain stem are reduced but the errors in the dentate gyrus and cerebellum are significant (arrows in E). The right hand column shows the result of ANTs (Avants et al. 2008) with affine and SyN and default settings. There is a reasonable overlap in the dentate gyrus but significant mismatch in the cerebellum and olfactory bulb (arrows in F).

### 2.4 Quantitative loss function

The goal of registration is to obtain the optimal composite transformation T: that can transform the moving image volume (M) to the same space as the fixed image volume (F). For a single transform stage, the transformation can be obtained from optimizing the loss function: in which S is the delineation of the similarity between F and transformed M. Common similarity evaluation metrics include mutual information (MI), cross correlation (CC), mean square error (MSE), which capture how well the two images are matched based on the joint histogram or signal intensities. Since we may use these metrics during registration, using the same metric repetitively for evaluation is unacceptable. At the same time, MSE, CC, global MI etc. by their intensity-based or histogram-based principles will not generate a stable predictability map between cross-modality datasets LSM and MRH. Therefore, we need to devise our own loss function.

The initialized LSM data is warped to MRH space with a combination of registration steps from ANTs. Our workflow encompasses multiple types of registration, and each type has different settings of similarity metrics for optimization and multi-resolution coarse-to-fine refinement. The loss function should evaluate the cumulative consequences of these steps. We devised a loss function based on a large group (~200) of fiducials. These fiducials were generated by an experienced researcher and consisted of matched pairs of points in MRH and LSM. Assuming the composite transform generated from our workflow is T, applying T to the LSM transforms these fiducials to the MRH space. The distance between one MRH fiducial (r_mr_) and its corresponding transformed LSM fiducial (r_lst_) is regarded as displacement from ground truth, and the average displacement i.e. L2 norm is used as the loss score, i.e.

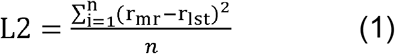

### 2.5 Optimization and validation

The registration transform can be separated into the category of linear and non-linear transforms. To reduce the computation, a complicated registration task should start from the linear transforms to adjust the position, orientation, and scaling of the moving image to coarsely and globally match the fixed and moving images, represented by the affine transformation equation:

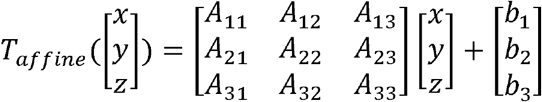

Then, application of non-linear transforms will deform the grid to locally match the fine details of fixed and moving images. From the popular options of non-linear transforms, we choose B-spline and symmetric diffeomorphic normalization (SyN) registration methods based on their efficiency on large, complicated datasets.

B-spline relies on the controls points to adjust local transform until reaching the minima of the loss function: for one b-spline transformation along x axis with n+1 control points {*P*_0_, *P*_1_,…, *P_n_*} and k degrees of freedom,

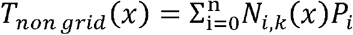

in which *N_i,k_* are the basis functions defined by the Cox-de Boor recursion formula and *X* = {*x*_0_, *x*_1_,…, *x*_*n*+*k*+1_} are the knots sectioned within the range [*x*_0_, *x*_*n*+k+1_] which x lies between.

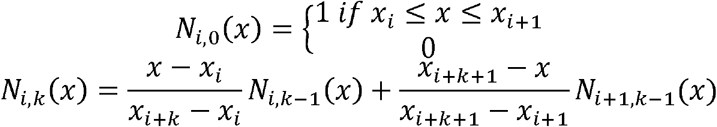

The curve defined by B-spline is a conjunction of multiple polynomial curves which only depends on a local group of k+1 control points. Based on the basis functions and the zero-order parametric continuity of B-spline, changing one control point will only influence the local neighborhood on the grid instead of propagating further. Therefore, B-spline can generate complicated and localized deformations flexibly and is computationally efficient when dealing with many control points.

SyN, as a representation of diffeomorphic algorithms, generates pixel-wise transformation based on symmetrical and invertible displacements and velocity fields. SyN is implemented on the Insight ToolKit platform and based on Large Deformation Diffeomorphic Metric Matching (LDDMM) principles. As an improvement, it develops the symmetry between the fixed and moving images, i.e. instead of maximizing the similarity between *T* · *M* and F, SyN maximizes the similarity between *φ*_1_(*m*, *t*)*M* and *φ*_2_(*f*, 1 – *t*)*F*, in which *t* ∈ [0,1], m and f are the respective identity positions of M and F, and *φ*_1_, *φ*_2_ are the respective correspondence maps from M to F, and from F to M. Based on the backward and forward symmetry, t=0.5. The optimization problem is then based on the equation:

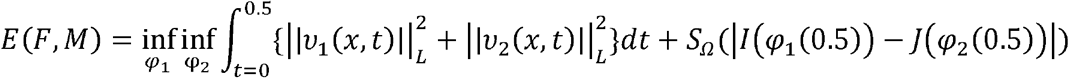

to minimize both the pixel displacement and the difference between *I*(*φ*_1_(0.5)) and *J*(*φ*_2_(0.5)), in which *ν*_1_ and *ν*_2_ are velocity fields in the opposite directions, *S_Ω_* is the similarity measure across the whole (x,y) surface. The advantage of SyN is the low computational cost and the preservation of the image topology.

Other than the transformation types, another factor influencing the registration is the selection of similarity metric. The most common similarity metrics include cross correlation (CC) and mutual information (MI).

CC defined by

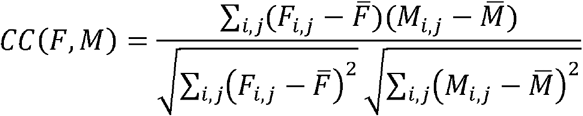

will reach the maximum when the intensity values in two images are related with a linear relationship. MI defined by:

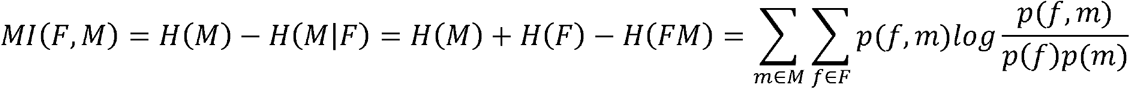

originates from information theory and measures how much information of one image can be predicted correctly from another image which is already known. MI focuses more on the global similarity between the histogram distribution of two images.

We call each individual transform together with a similarity measurement metric as a stage. Beside the transform types and similarity metrics, the stages usually have many parameters which increases the complexity and duration of optimizing the transformation. In each stage, we employ multiresolution method, which initially performs the registration at a lower resolution with fewer control points and then samples the control points to a higher resolution in the following levels, which will efficiently converge the loss function without consuming much computing resources.

The optimization was initially performed on data that had been downsampled to 45 μm to allow a broad search of parameters. This was followed by a refined optimization with both MRH and LSM data at 15 μm voxel size for parameter refinement. The first stage of optimization was conducted using diffusion weighted (DWI) MRH and LSM stained with a generalized cell marker (Syto16). As the workflow might encompass multiple registration stages with different transforms and similarity metrics, and each stage encompasses multiple parameters, it is uncertain which combination of registration settings can provide the best performance. Therefore, the workflow is optimized with an inverse pyramid optimization pipeline, i.e., starts from the large structures and similarity metrics combination, and proceeds to the parameters in specific stages respectively. Data transformed with the optimized settings for one stage of the pipeline served as the input for tuning the next step in the pyramid.

The optimization pyramid (Table 2) includes:

- Stage 1 focuses on optimizing large structural details. This step employs linear registration (rigid and affine) followed by non-linear registration (b-spline syn and syn). Each stage uses the same default parameters. In stage 1, P1_02 i.e. Affine (Default) + B-spline Syn (Default) + Syn (Default) yielded the lowest loss score so its output served as the input for stage 2.
- Stage 2 focuses on similarity metrics, i.e., mutual information or cross correlation.
- Stage 3 adjusts the multi-resolution settings with different number of multi-resolution layers, shrink factors (i.e., down-sampling) and smoothing sigmas (i.e., the radius of Gaussian filter).
- Stage 4 focuses on spline distance, a specific parameter in b-spline syn.
- Stage 5 alters the multi-resolution settings with different number of multi-resolution layers, shrink factors and smoothing sigmas.

**Table 2.**
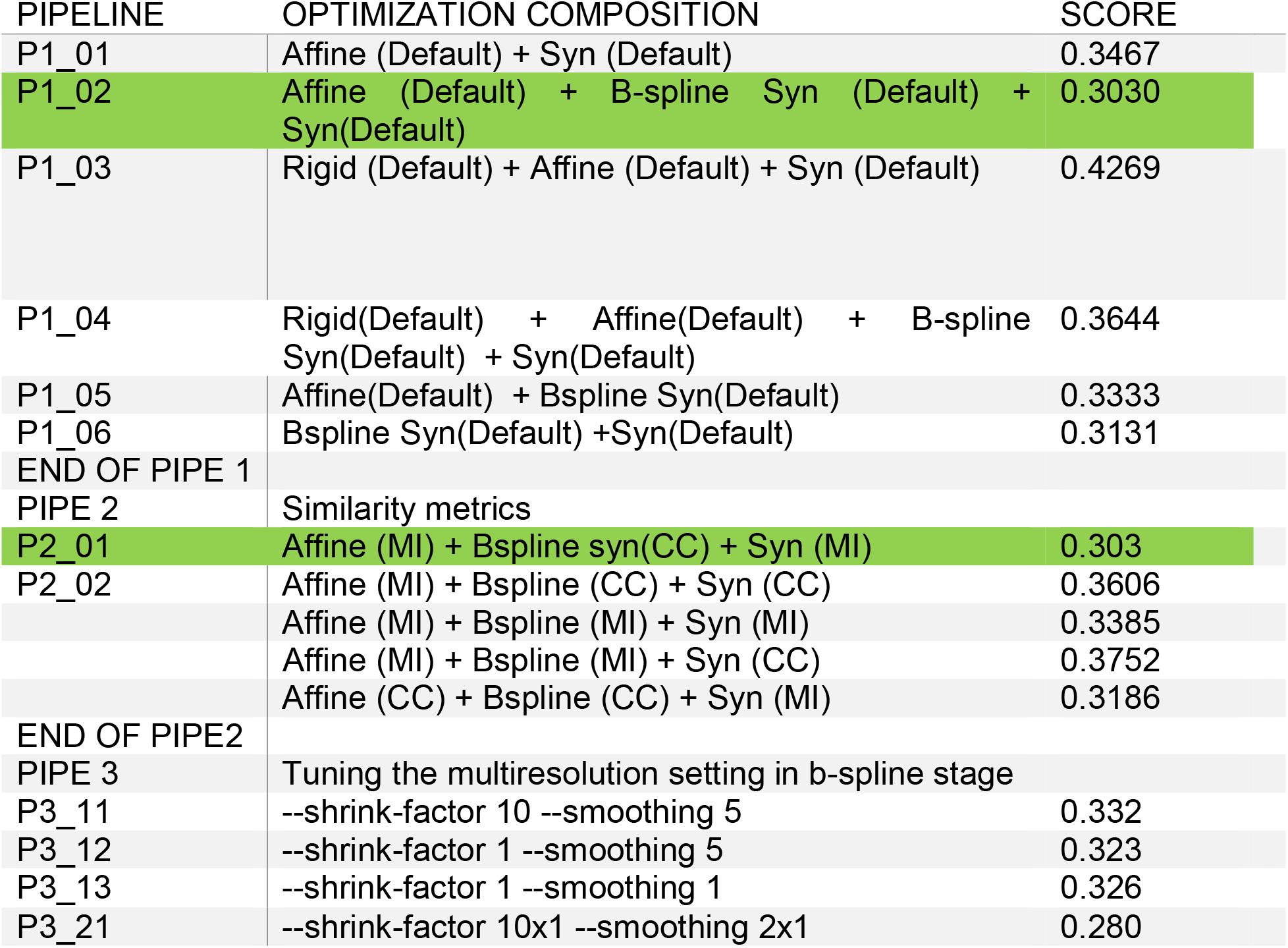
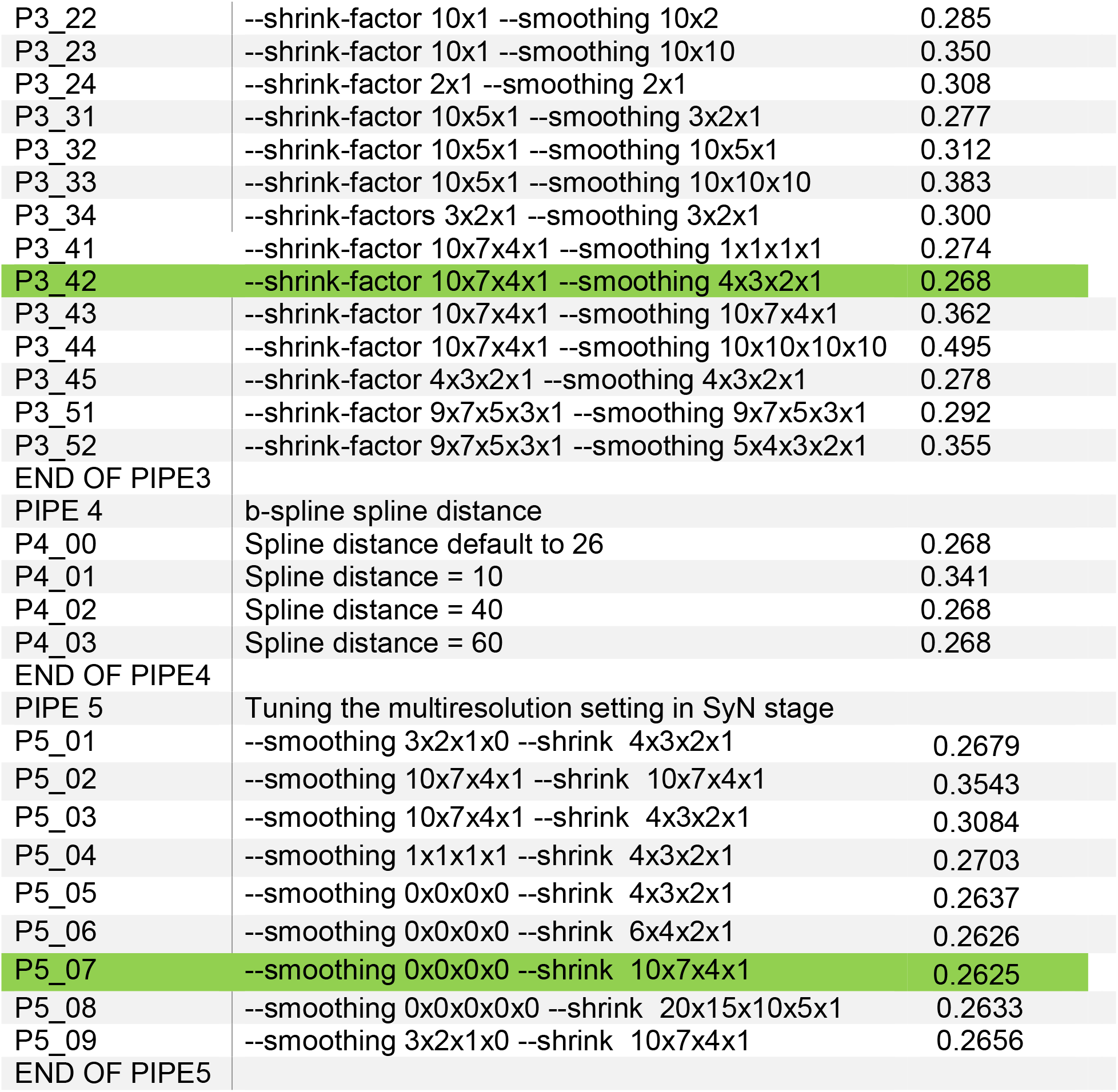
Pipeline initialization @ 45 μm resolution. The steps and parameters for the pipelines that were tested are summarized.

**Table 3.** Optimization of pipeline @ 15 μm resolution.

| Pipeline | Composition | Score | Time |
| --- | --- | --- | --- |
| P6_01_H | --shrink-factor 10x7x4x1 | 0.2419 | 3d17h |
| P6_02_H | --shrink-factor 30x21x12x1 | 0.2834 | 6d12h |
| P6_03_H | Finer affine<br>--shrink-factor 30x21x12x3 | 0.2147 | 2h29m |
| P6_04_H | Finer affine<br>--shrink-factor 30x21x12x1 | 0.2544 | 6d12h |
| P6_05_H | --shrink-factor 40x28x16x4 | 0.2232 | 2h43min |
| P6_06_H | --shrink-factor 20x14x8x2 | 0.2314 | 18h12m |
| P6_07_H | --shrink-factor 10x7x4x2 | 0.2382 | 10h29m |

The optimization starts from the arrangements of different stages with the same parameters based on the default setting (from ANTS python https://antspy.readthedocs.io/en/latest/registration.html), which includes 4 rounds of registration iterations on different resolution.

The pipeline with the lowest loss score is labelled in green at each stage. The starting point for all the comparisons is the initialization using ~20 manual landmarks (Figure 3B,C). The comparison between a pipeline that is less accurate (p2_02, L2 =.436), and the optimal pipeline @ 45 μm e.g. (p3_42, 0.268) is shown in Figure 3C,G and D,F. The improvement is evident. But extending the pipelines to 15 μm data must consider computational time coupled to careful inspection of the results. This last stage of optimization is more nuanced depending on computation time and the combination of LSM/MRH contrasts (e.g., DWI/Syto, FA/NeuN) which is discussed in more detail in Section 3.2The optimization @ 15 μm is started from pipeline p6_01, which has the same registration setting with the optimal pipeline @ 45 μm (P3_042). However, it does not generate the expected improvement, it overfits the data, and is 27 times slower. Optimization @ 15 um is not efficient because the data sets are 27 times larger than the 45 μm. Therefore, the final optimization takes both the computation time and quantitative score as assessment of a pipeline (Table 2). Comparison between the best pipeline at 45 μm (P3_042) and a pipeline p6_06H optimized on 15 μm is shown in Figures 3D,H and Figure E,I

**Figures 3.**
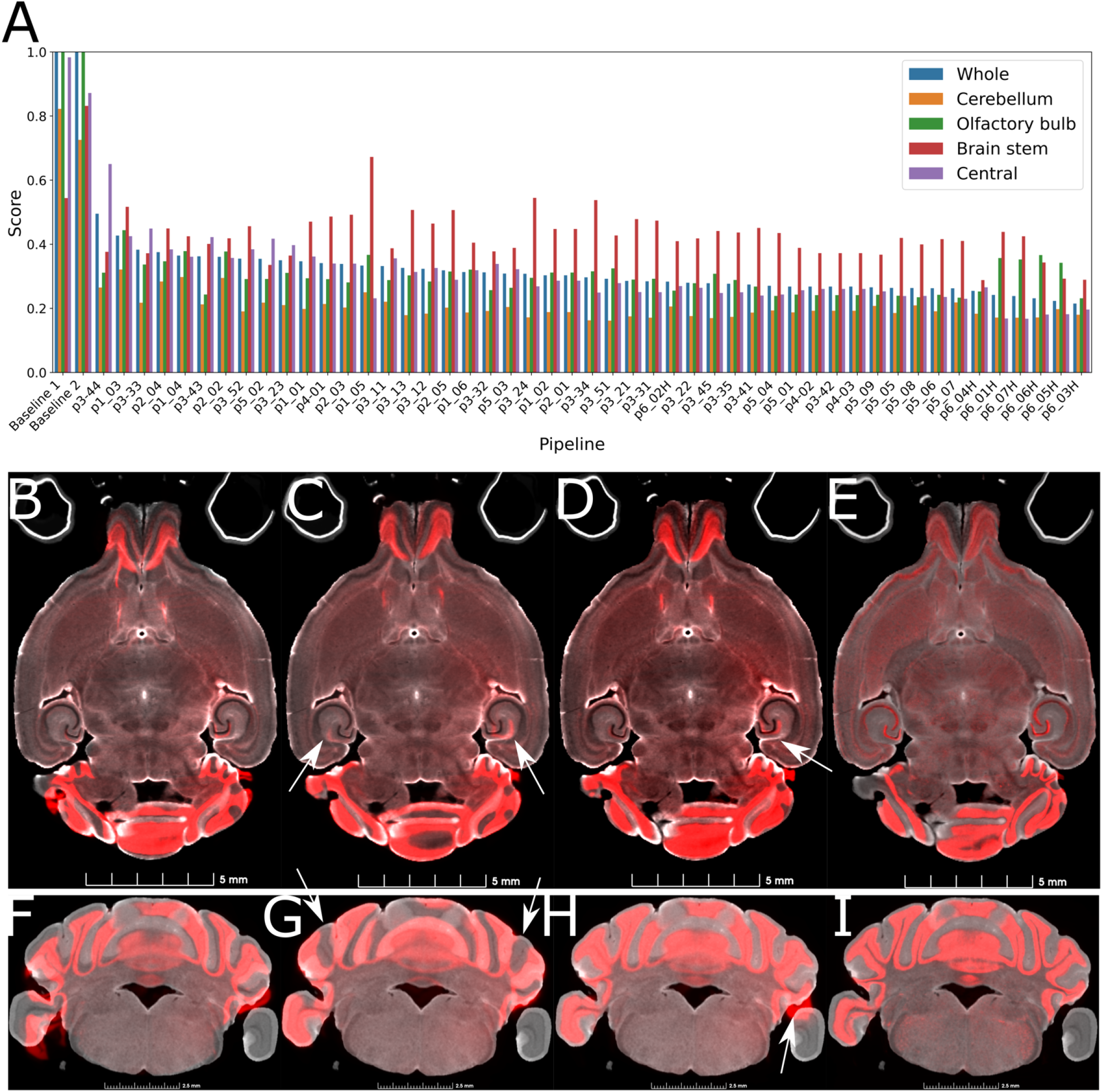
demonstration of the range of results derived from the varied pipelines. Figure 3A plots the L2 norm for the pipelines listed in Tables 2 and 3 registering Syto16 to DWI (Specimen 191209). In figure 3B,F, due to the sparsity of the initialization landmarks, the initialization procedure prepares the registration starting point well in the central slice of the brain (figure 3B) but not in non-central slices e.g. figure 3F. Figures 3C,G, and D,H show results at 45 μm with L2 norms of .361 and .268 respectively. In figure 3C,G, pipeline p2_02, there are significant errors in the cerebellum even with the strong coherence between the LSM and DWI in the cerebellar cortex. Figure 3D,H with pipeline p3_42 performs better. This example was chosen to highlight one of the frequent challenges in the LSM data. The parafloculoss is missing on the right side of the brain. This is a common problem arising when scanning tissue that has been removed from the skull. The broken symmetry in the data gives rise to the potential of generating false data. Note for example, in the axial image in Figure 3C the asymmetry in the alignment of the dentate gyrus in the central part of the brain. Finally, a comparison of Figure 3D,H (@ 45 μm) and Figure 3E,I (@ 15 μm) with pipeline p6_07_H demonstrates the utility of performing the registration using the higher resolution data.

## 3. Results

### 3.1 Optimization of pipelines

Figure 3 A shows registration results for all pipelines listed in Table 2 rank ordered according to the L2 norm for the whole brain. The L2 norm is also shown separately for the cerebellum (CB), olfactory bulb (OB), central section of the brain (C) and brain stem (BS). Each region poses unique challenges to the algorithm. The contrast is very high between the white matter and intensely staining granular cell layer in the cerebellum in both the NeuN and Syto images. And there is comparable strong contrast in the DWI. Thus, the L2 norm for this cerebellar region converges to a low value for all the pipelines. In the central part of the brain, the dentate gyrus, fimbria, and corpus callosum all provide unambiguous landmarks and fine tuning the pipeline leads to gradual improvement in the score. The olfactory bulb shows a similar effect, but the score does not converge to as low a value. This may be because the olfactory bulb is one of the most distorted regions of the brain. And there are frequent tissue tears (arrow 1 in Figure S1). Finally, the brain stem is the most challenging region for registration as evidenced by high L2 norm and the high variability between different pipelines. The cause of this is again evident on inspection of the sagittal LSM and MRH imaged in Figure 1 C and Figure 2 D,F. The spinal cord in the LSM is grossly misplaced from its natural position forcing the algorithm into large displacements.

The transform obtained from the 15 μm registration is applied to the full resolution LSM data through the python interface of 3D Slicer, in the order of their generation. The time to apply transforms to full resolution LSM data (~300GB) is ~2hours. The Python and bash codes for running ANTs, obtaining quantitative evaluation and visualization, applying transforms and warps in Slicer are available upon request.

### 3.2 Impact of image contrast

The registration success depends on the similarity between the information presented by the fixed and moving volumes. The initial work described above varied the pipelines while registering Syto16 to DWI. This section of the manuscript uses a fixed pipeline (p6_07H) while varying the combinations of LSM/MRH images. The DTI pipeline produces 11 different scalar images, each highlighting different tissue architecture. And the anatomic landmarks in the LSM vary widely depending on the immunohistochemistry used. There are an enormous number of combinations. Figures 4A-F show representative comparisons derived from specimen 200316. The auto fluorescent image (Figure 4A) is frequently used to drive registration to the auto fluorescent image in the ABA. NeuN (Figure 4B) and MBP (Figure 4C) are of particular interest to our work in aging. The DWI (Figure 4D) is created by averaging all the (registered) diffusion weighted images producing high contrast to noise with many anatomic landmarks throughout the volume. Cortical layer definition and contrast in the dentate gyrus are particularly high in this volume. There are strong similarities between NeuN (Figure 4B) and DWI (Figure 4D). The FA image (Figure 4D) is a logical choice as it highlights white matter. And the RD image (Figure 4F) is a putative marker of myelin integrity that might map well to the MBP. This specimen has the largest number of fiducials providing the largest global measure of alignment.

**Figure 4.**
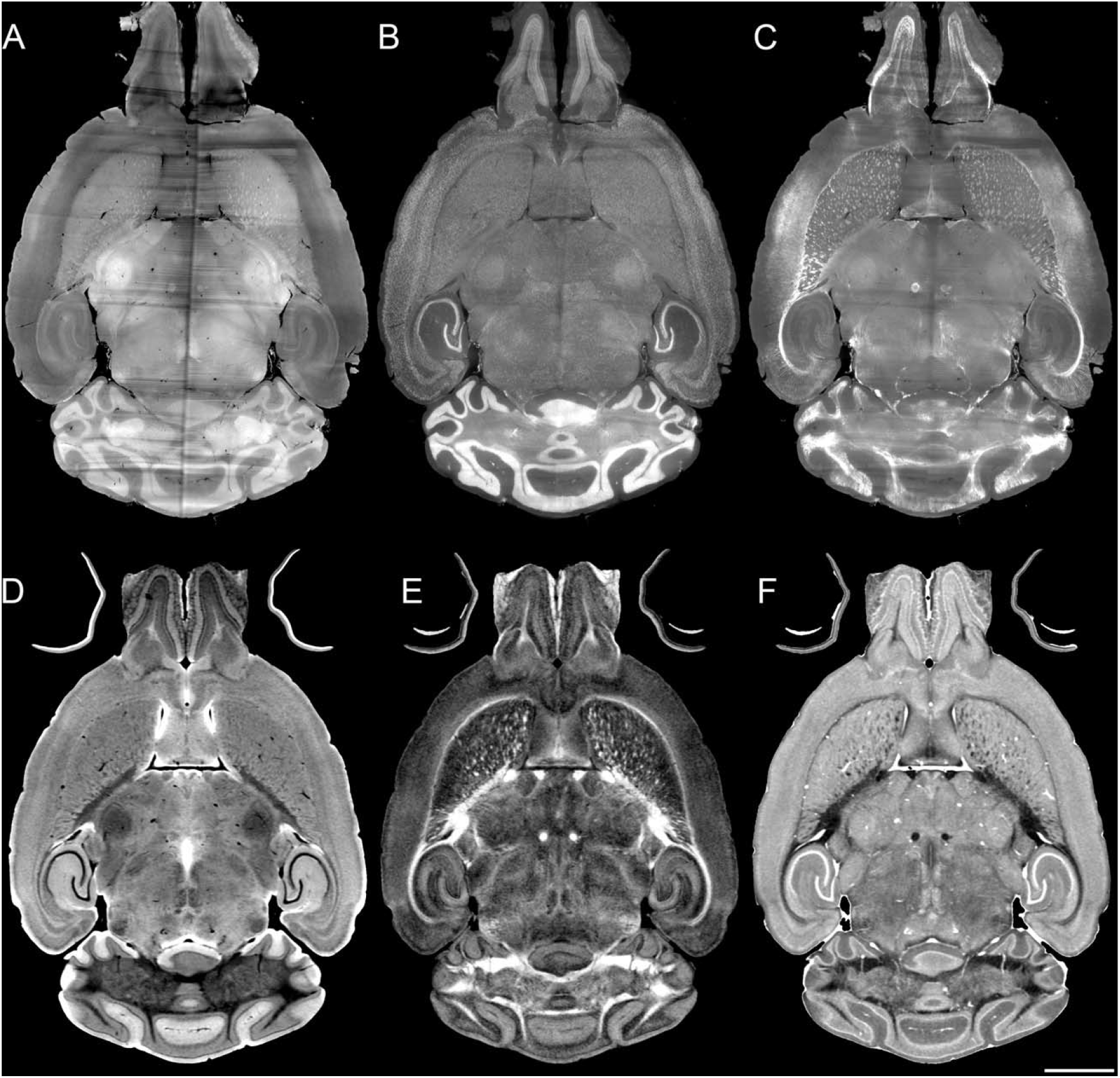
LSM of different stains and MRH of different contrasts. A) Auto fluorescent B) NeuN C) MBP D) DWI E) FA F) RD Scale bar is 2 mm. (Specimen 200316)

Two pipelines were chosen for more careful comparison: p6_03H and p6_07H. Because of the similarities between NeuN and DWI, this combination was chosen to evaluate the two pipelines in three different specimens. Table S1 and Figure S1 in the supplement summarize the comparison. P6_03H is faster than p6_07H and for one specimen (191209) yielded a lower L2 norm. The resulting volumes were imported into Imaris to allow interactive review of the relative success of the registration across the entire volume. Figure S1 demonstrates that p6_03H results in consistent subtle misregistration in the dentate gyrus that is absent in p7_07H.

Table S2 summarizes an exhaustive comparison of p6_07H across five specimens with 15 different pairs of images. Specimen 200316 with the largest number (200) of fiducials was run twice with different initializations. Specimens 190108 and 191209 are from the BXD series providing a strain with different anatomy than the B6. Comparison of the L2 norms between specimens is not appropriate since each specimen has a different set of fiducials. This highlights some of the limitations in using fiducials as a quantitative metric for comparison of the quality of a registration. The precision of fiducial pairs will be biased by the reader placing the pairs. This results in a lower (nonzero) level which will vary between specimens that is dependent on the reader/fiducial e.g., an average error of 135 μm for the NeuN/DWI combination for specimen 200803 with 51 fiducials and 235 μm for specimen 191209 with 175 fiducials. However, comparison of the L2 norms across the different registration combinations within a specimen can provide useful insight into which pairs provide the best registration. For example, mapping MBP to RD is one of the least successful combinations. Mapping NeuN to DWI or Syto to DWI yields one of the lower L2 norms for all the specimens. The duplicate comparison for specimen 200316 highlights the stochastic nature of the registration with a 12% difference in the L2 norm (NeuN+DWI) between the two runs. But the relative scores of varied combinations of mapping remain unchanged.

One of the more surprising results is the success of the auto fluorescent/DWI combination. Figure S2 shows the results of registration using p6_07h with two image combinations: Autofluorescence (autoF) to DWI and NeuN to DWI with specimen 200316. The transforms generated with the autoF to DWI registration was then applied to the NeuN. The registered pairs (NeuN to DWI) for both transforms were interactively reviewed in Imaris to discern areas in which the transforms differed. The target image (DWI) is displayed in yellow, and the moving image (NeuN) is displayed in green. In Figure S2A (NeuN to DWI) there are subtle errors in alignment in the cerebellum that are not evident in the autoF/DWI pair. Yet the internal structures e.g., the dentate gyrus seem to be comparable. Comparison of the moving images C) NeuN or D) autoF, highlight the high contrast granular layer in the NeuN image and the relatively flat contrast in the auto fluorescent image. The high contrast in this granular layer dominates the registration since the NeuN stain in the outer edge of the brain is nonexistent. Registration using the auto F is more successful since the contrast in the cerebellum is quite flat. This highlights one of the most challenging aspects of this task i.e., the registration of two volumes with completely different sources of contrast.

Landmark comparison of the NeuN to DWI registration was undertaken to gauge the quality of registration away from the edges. The NeuN and RD images were overlayed in Imaris. The strong vascular signal from the RD was helpful particularly in identifying the plane of section. Vessels were identified in the NeuN (Figure S3) and magnified. The plane of the vessel cross section was adjusted until the bifurcation was evident and a fiducial was marked. The same steps were performed with the RD. The process was executed on 11 different vessels spread throughout the brain. The mean displacement of the 2 images was 22 ±14 μm.

### 3.3 Volume corrections to LSM (190108)

The most common way of delineating brain regions on an LSM image is via registration to an atlas (Tappan et al. 2019) (Perens et al. 2021) (Kutten et al. 2016) or registration of the atlas to the volume under study (Goubran et al. 2019). The most commonly used atlas is the ABA i.e. the CCF v3 3D template constructed from a population of 1675 young adult B6 brains using autoF (Wang et al. 2020). We have created an MRH atlas with a subset of these labels using specimens 200302 (male) and 200316 (female) (Johnson et al. 2022). In Figure 5, we used this MRH atlas to estimate the roegional volume changes in the LSM images from tissue swelling in specimen 190108. This specimen (111 day BXD 89) is representative of our broader interest-understanding the genetic basis for age related changes in the BXD family (Ashbrook et al. 2021). We registered the NeuN to DWI for specimen 190108 using pipeline p6_07H. Labels were registered to the DWI of specimen 19018 from our reference B6 atlas (200302) using our MRH registration pipeline (Anderson et al. 2019). The transform that was generated was inverted to transform the labels on the DWI back to the uncorrected NeuN volume. Figure 5 A,B shows the NeuN volume before and after correction respectively. Note the changes in the width is larger than the change in length highlighting the nonuniform distortion. This is even more apparent in Figure 5 C and D which shows a sagittal cross section before and after correction.

**Figure 5.**
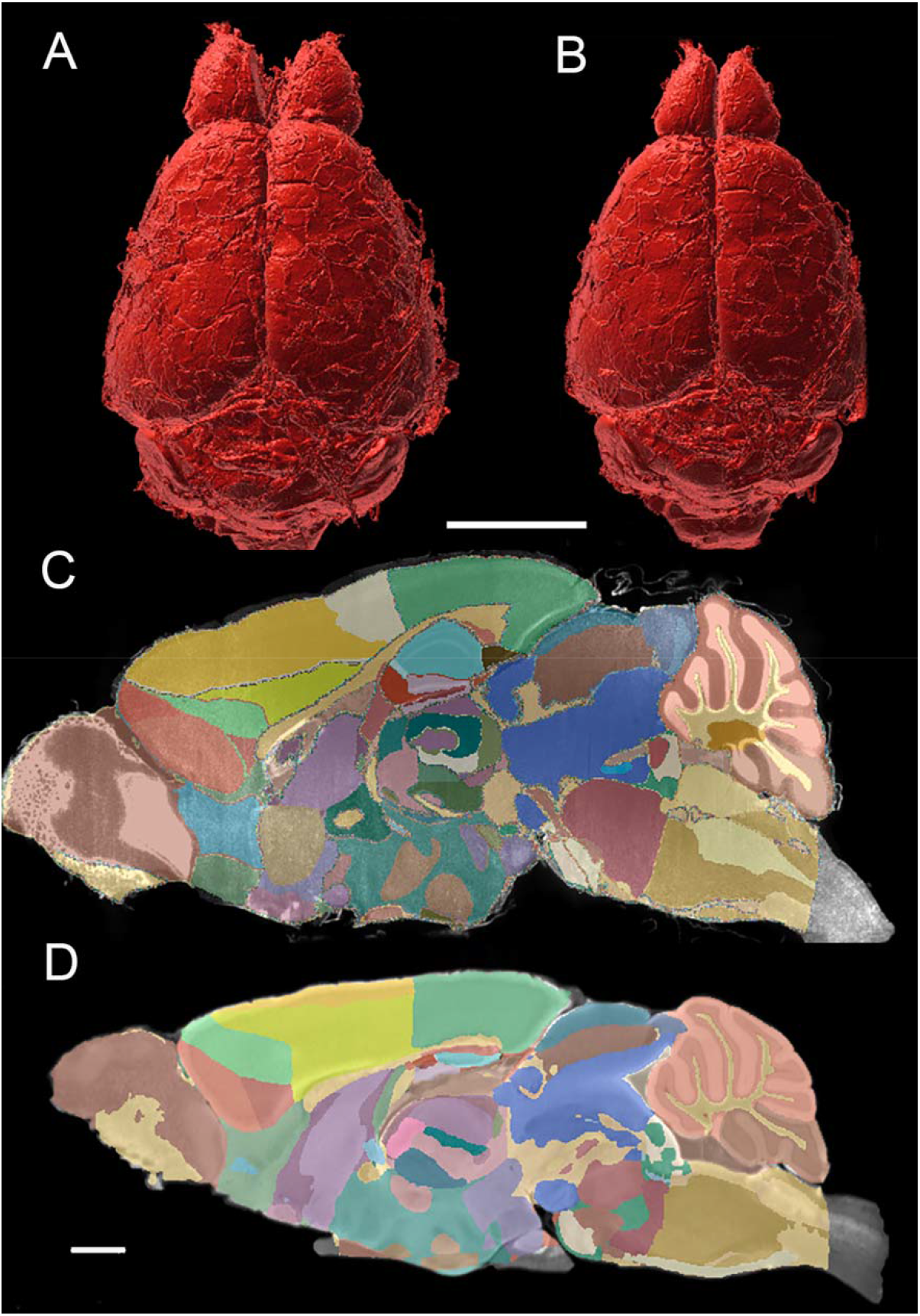
Distortion correction of the LSM data by registration to the MRI of the same specimen (190108). (A) Surface rendering of uncorrected LSM volume and B) corrected LSM volume: scale bar is 5 mm. C and D) midsagittal section of the labels on the LSM data before C) and after D) correction: the scale bar in C,D is 1 mm. The distortion is present both within the plane of section and across the plane making it difficult to define identical planes. The highlighted edges in C are an interpolation artefact

Figure 6 summarizes the change in volume for the 50 largest regions of interest. The abbreviations are consistent with those used by the Common Coordinate Framework (CCFv3). We have created a reduced set of labels (rCCFv3) by combining some smaller structures into a single label since mapping the smaller structures is more highly variable. A summary of those labels can be found in (Johnson et al. 2022). The magnitude and variability are significant. The olfactory bulb (OB) is nearly 80% larger in the uncorrected data while the corpus callosum (cc) is ~10% smaller. The problem is compounded when comparing specimens as the differential swelling varies. And it varies considerably between different clearing methods. These variations must impact the shape of the structures. Figure S4 demonstrates the impact on the non-uniform distortion on the hippocampus, a region of particular interest in age related neurodegeneration (Sabuncu et al. 2011) (Katabathula, Wang, and Xu 2021).

**Figure 6.**
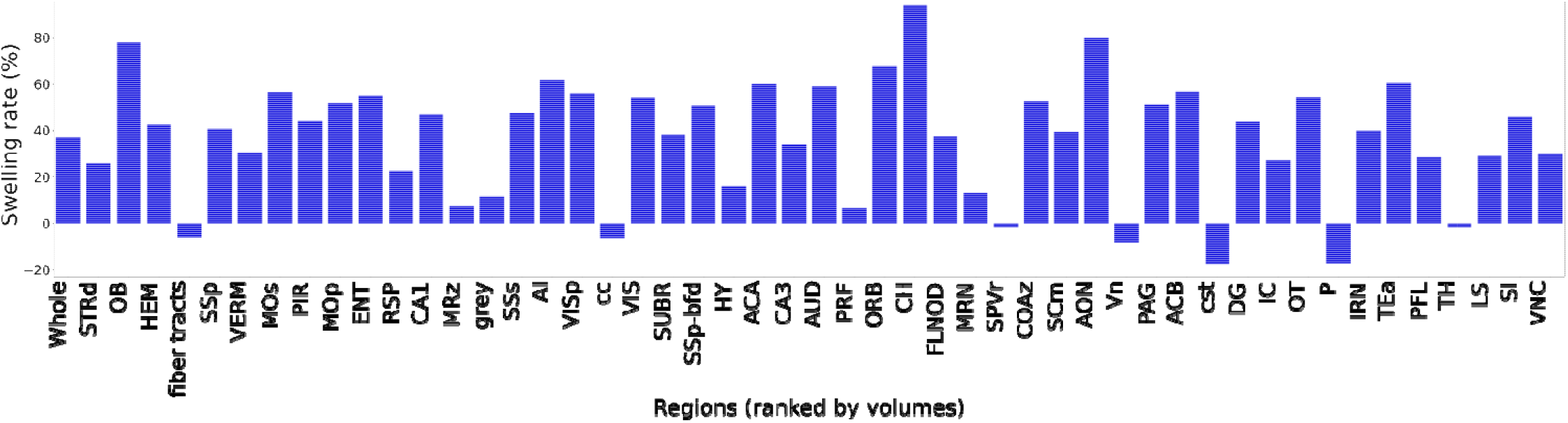
Bar plot of ratio of the volume before and after registration. The regions are ranked by the ROI volume. The rate is obtained by—————.

## 4. Discussion

Our main initiative originates from the fact that there are resourceful neuroscience insights from the unique morphology of every brain. Both the shapes and regional quantitative data provide essential information for neurogenesis research, but the necessary step of segmenting brain regions commonly relies on registering the brain images to some certain atlases, which obscure the unique information of brains from different genetic strains and ages, etc. In this paper, we developed a method to register the cellular brain images to MRH, which maintains the ground-truth of brain morphology inside the skull. In the space of MRH, the region segmentation (rCCF labels) is applied on cellular brain images, which can maintain the brain morphology in the skull.

To register LSM to MRH with little errors and efforts, we conquered several challenges, i.e., correcting the significant and irregular distortion in the LSM between the two very different contrast modalities, with LSM data also large in the size per channel per specimen (~500GB). Common popular algorithms can hardly register the data successfully, mostly due to the significant distortion in LSM. We have solved the registration problem with an initialization involving ~ 20 landmarks and have optimized a combination of layers of different transformations and metrics to minimize a user customized L2 norm score.

From the optimization, we selected the registration workflow with a combined consideration on accuracy and time. The workflow (pipeline p6_07) we selected to process 190108, for example, takes an average of 7.5 hours in registration, with the L2 norm score for this registration at 135 um. The workflow shows robustness in multiple specimens. Our approach takes advantage of the high spatial and contrast resolution in the MRH images to provide internal landmarks the drive the registration locally across the whole brain which is evident from the small mean displacement (~22um) of fiducials, which are picked at the junctures of vessels in both MRH and LSM.

As both MRH and LSM include varied contrasts (Figure 4), we did experiments to find the best combination of different diffusion scalar images and immunohistochemistry with LSM. A surprising conclusion is that registrations between dwi and autofluorescence or NeuN are similarly good. Our explanation is, the flatness of autoF might benefit registration by driving it with its solid contour, while the registration between NeuN and dwi is driven by the multiple internal landmarks. The practical consequence for our use is that we will not have to acquire an autofluorescence image freeing up a channel in the LSM for a more useful cytoarchitectural measure i.e., NeuN.

Multiple groups have developed methods for automated labeling of 3D optical images from cleared specimens (Perens et al. 2021; Kutten et al. 2016). These approaches rely on the Allen Brain Atlas as the reference (Wang et al. 2020). We are interested in mapping the age-related changes across multiple strains (both genders). Registration of these data to the young adult male C57 that is the core of the ABA would obscure most of the morphologic changes of interest. Reiner et al used MRI of a fixed mouse brain to measure the degree of distortion from tissue processing with iDisco but their MRH images were of a half brain taken with a relatively low contrast gradient echo out of the skull. Labeling relied on mapping the autofluorescence image to the ABA. The MRI was not used in this step. Gourban et al have developed a pipeline that is similar to that which we report here(Goubran et al. 2019). Our work differs from their approach in four ways. Our dMRI protocols acquire data @ 15 μm vs 200 μm i.e., a difference in voxel volume of 2370 X with the commensurate challenge of larger image arrays. As demonstrated in Figure 3E,I registration with the full resolution MRH (15 μm) makes a difference. We hope to explore a range of immunohistochemistry stains. Table S2 provides an excellent starting point for evaluation of many of the alternatives. Finally, our pipeline takes advantage of a truly isotropic 3D MRH atlas of the brain in the skull to which rCCF3 labels have been matched. Our approach provides an efficient method for segmenting brain regions in LSM data mapped in the MRH space of the same specimen which will allow quantitative study of cytoarchitecture e.g., cell density along with connectivity. The contrast study also would be a fruitful area for the further work. For example, a broader study could consider synthesizing synthetic contrast from combinations of scalar dMRI images that might contain complementary information or using machine learning to transferring the contrast from LSM to MRH to reduce the registration difficulty due to different contrast distributions. Artificial intelligence may well provide new avenues to improve the registration quality and efficiency.

## Acknowledgements

We are grateful to Leonard White, Ph.D. in the Duke Department of Neurobiology and Robert W. Williams, Ph.D. Univ of Tennessee Health Science Center Chair of Genetics for helpful guidance in neuroanatomy. We thank Tatiana Johnson for technical assistance in preparing the manuscript. This work was supported by National Institute of Aging R01AG070913 (to GAJ and RWW)

