## Supplemental Figures &Tables for "Restoring morphology of light sheet microscopy data based on magnetic resonance histology"

**Supplementary Materials**

| Specimen num. | Fiducial pairs | p6_03H | p6_07H |
|---|---|---|---|
| 200316 | 200 | 0.1835 (2.4h) | 0.16(7.7h) |
| 191209 | 175 | 0.2147 (2.5h) | 0.2382 (10.5h) |
| 200803 | 50 | 0.2152 (3.8h) | 0.1351(9.5h) |

Table S1. Comparison of registration L2 norm and registration times for three different specimens (Syto16 to DWI) using two of the more successful pipelines. P6_07H is three times slower than p6_03H, but it has a lower L2 norm for two of the three runs. Examination of the resulting registrations (below) reveals subtle errors when using p6_03H that do not appear in p6_07H.

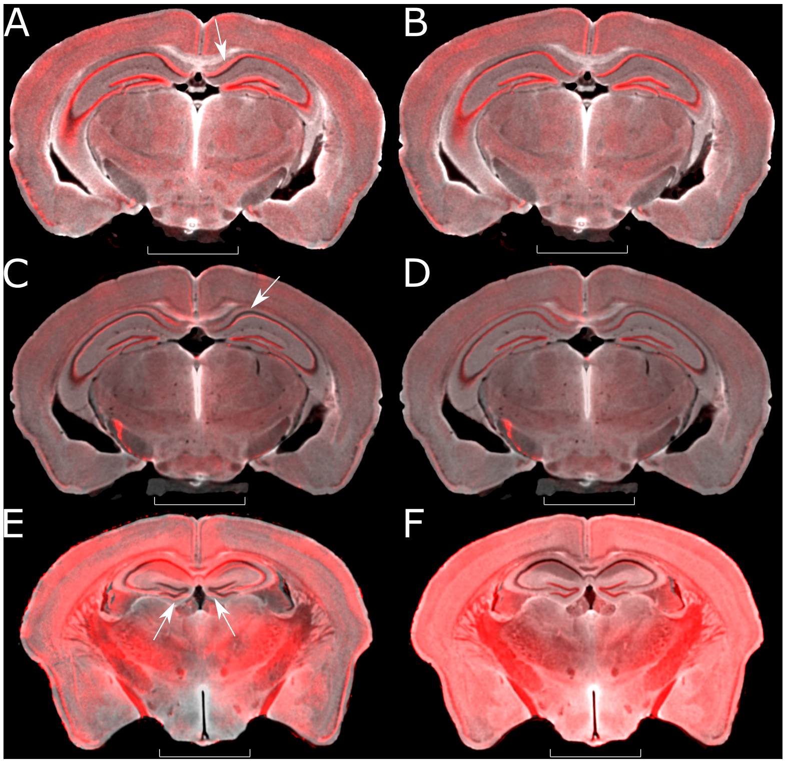


Figure S1. The performances of p6_03H and p6_07H in multiple datasets are different. The registered Syto (red) is overlaid on DWI (grey). In the first specimen (A and B), Syto registered to DWI by p6_03H (A) has slight displacement from DWI in the dentate gyrus (white arrow), the same Syto image registered by p6_07H overlaps the DWI much more effectively (B). Similar results are seen in two other specimens (C and D, E and F). The scale bar is 5 mm.

| Contrast\Specimen | 200803 | 200316 (1) | 200316 (2) | 200802 | 190108 | 191209 |
|---|---|---|---|---|---|---|
| NeuN+dwi | 0.135 | 0.160 | 0.182 | 0.222 | 0.179 | 0.235 |
| NeuN+fa | 0.169 | 0.192 | 0.224 | 0.274 | 0.195 | 0.417 |

| | | | | | | |
|---|---|---|---|---|---|---|
| NeuN+rd | 0.147 | 0.164 | 0.194 | 0.247 | 0.191 | 0.224 |
| Syto+dwi | 0.137 | | | | 0.172 | 0.221 |
| Syto+fa | 0.1662 | | | | 0.2027 | 0.2305 |
| Syto+rd | 0.1369 | | | | 0.1843 | 0.2391 |
| autof+dwi | | 0.150 | 0.157 | 0.2184 | | |
| autof+fa | | 0.155 | 0.164 | 0.2391 | | |
| autof+rd | | 0.1554 | 0.1609 | 0.2465 | | |
| MBP+dwi | | 0.1684 | 0.1845 | 0.2372 | | 0.2347 |
| MBP+fa | | 0.165 | 0.1791 | 0.2517 | | 0.2578 |
| MBP+rd | | 0.1684 | 0.1846 | 0.2439 | | 0.3131 |
| IBA1+dwi | 0.16 | | | | 0.1727 | |
| IBA1+fa | 0.1565 | | | | 0.1883 | |
| IBA1+rd | 0.1652 | | | | 0.1903 | |

Table S3. Scores of different combinations of contrast/channels in multiple specimens. Specimen 200316 was processed 2 times.

| **Source Data** | **Algorithm** | **Name** | **Abbreviation** |
|---|---|---|---|
| DTI | ANTs | Average Baseline | Avg$b_0$ |
| | Average | Diffusion Weighted Image | DWI |
| | DTI | Mean Diffusivity | MD |
| | | Axial Diffusivity | AD |
| | | Radial Diffusivity | RB |
| | | Fractional Anisotropy | FA |
| | | Color Fractional Anisotropy | ClrFA |
| DTI | GQI | Isotropic Fraction | iso |
| | | Normalized Quantitative Anisotropy | nqa |
| | Calamante | Track Density Imaging | TDI |
| | | Color Track Density Imaging | ClrTDI |

Table S4. Summary of image types and abbreviations.

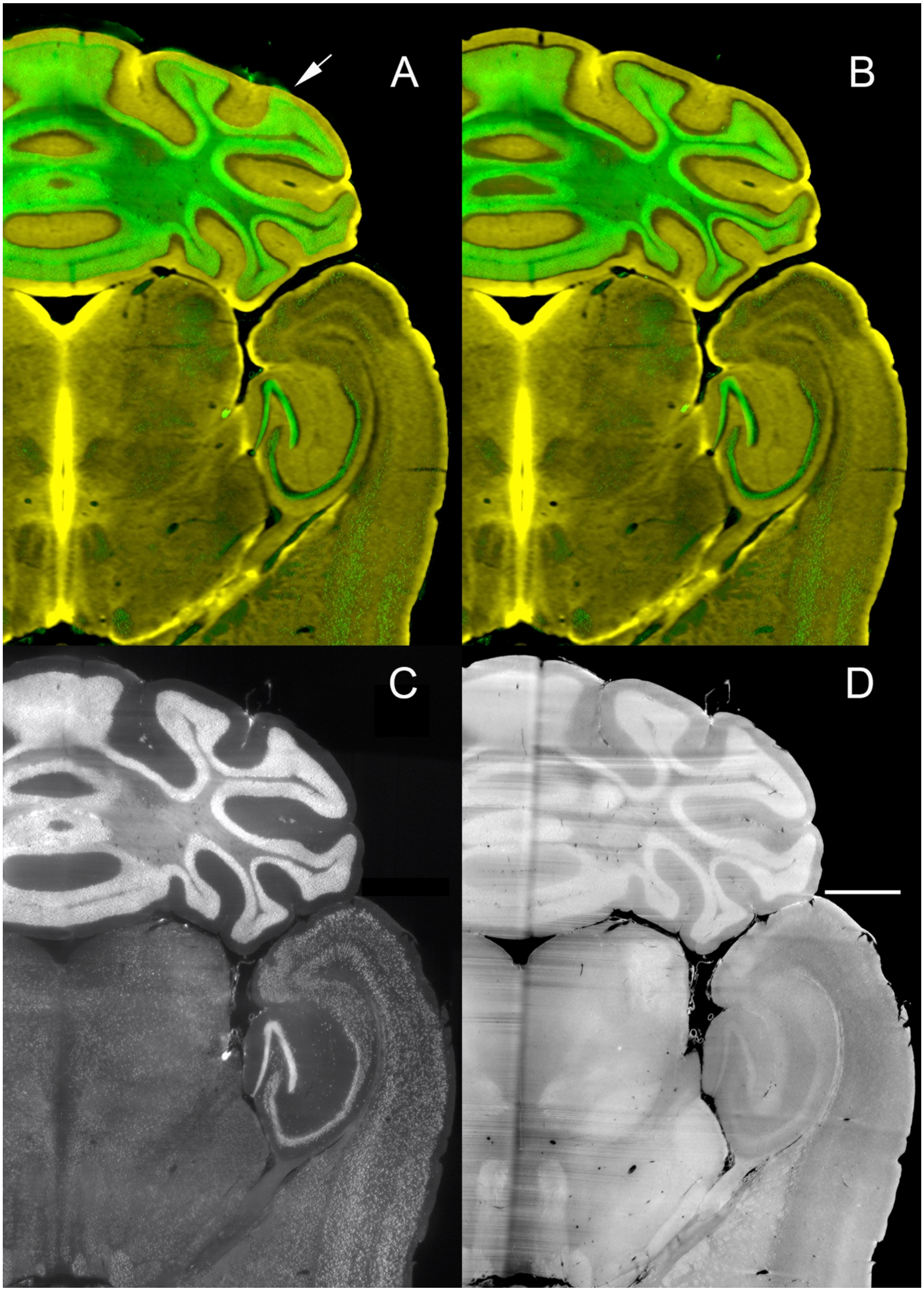


Figure S2. A) NeuN (green) aligned to DWI (yellow) using the NeuN as the moving image. B) NeuN (green) aligned to DWI (yellow) using the autofluorescence as the moving image. Note in

A (arrow) the misalignment at the periphery. Yet the internal structures eg dentate gyrus are quite comparable. The alignment error at the edge in A is ~ 150 μm. The scale bar is 1 mm.

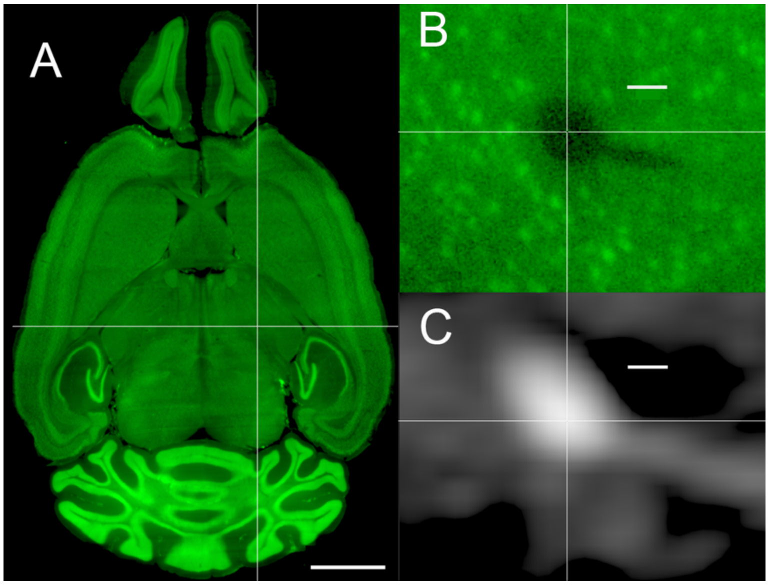


Figure S3. The selection of anatomically informed fiducials using vessels provides the most precise measure of alignment. We interactively select a vessel (Specimen 200316) in the NeuN image (cross hairs in A), magnify and adjust the location of the plane perpendicular to that which is displayed to find the plane where the branching vessel joins (B) and note the location in the 3D grid of the reference space in Imaris. The process is repeated in the RD image where the vessels appear bright. Using this method in 11 different points yielded a mean of 22±14 μm. The scale bar in A is 2 mm. The scale bar in B and C is 50 μm.

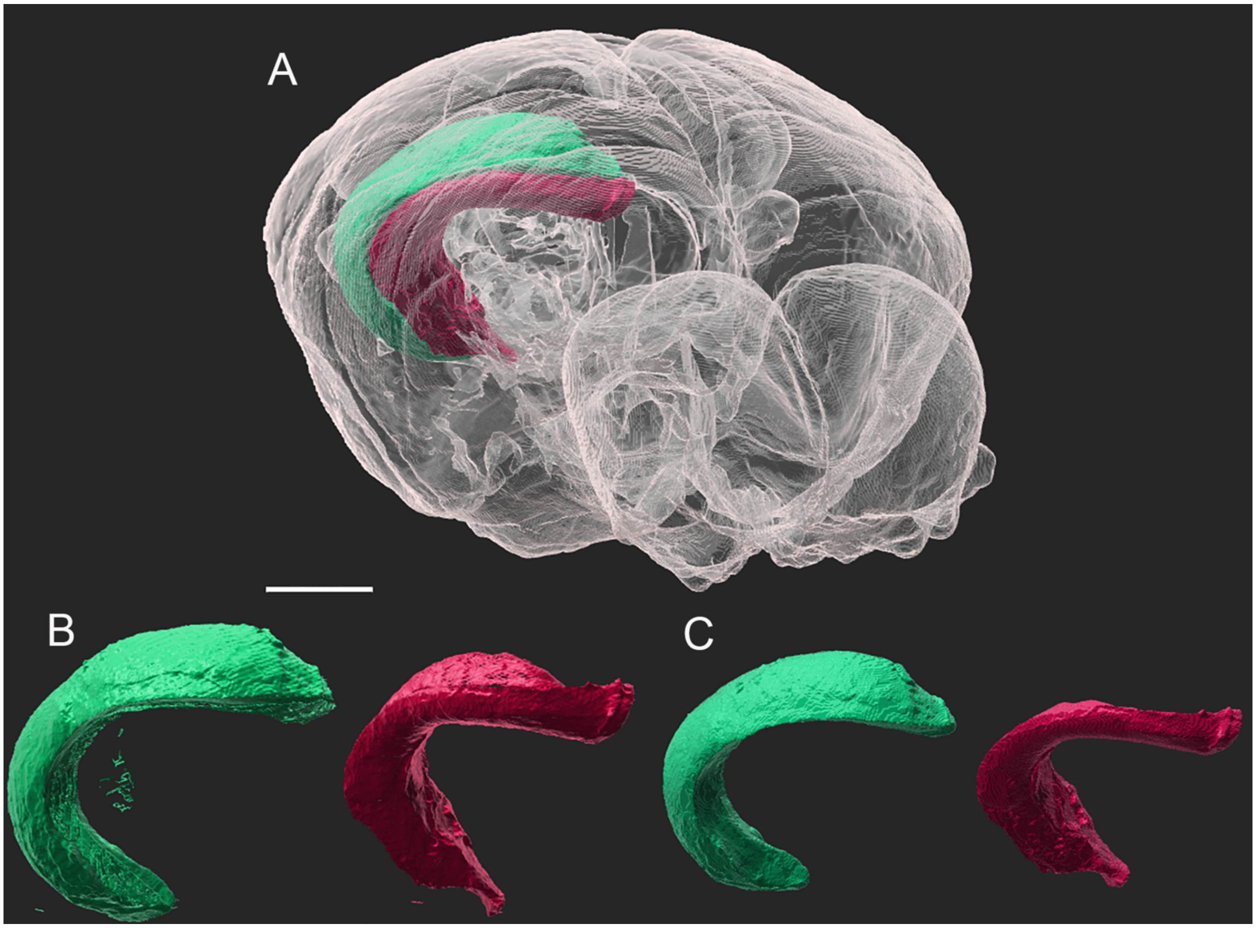


Figure S4. Volume rendered image showing hippocampal CA1 (green),CA3(red) A) in situ after registration and B) isolated for shape analysis before geometric correction; C) isolated for shape analysis after geometric correction. The scale bar is 2 mm.
